# Bayesian inference of differentially expressed transcripts and their abundance from multi-condition RNA-seq data

**DOI:** 10.1101/638817

**Authors:** Xi Chen

## Abstract

Deep sequencing of bulk RNA enables the differential expression analysis at transcript level. We develop a Bayesian approach to directly identify differentially expressed transcripts from RNA-seq data, which features a novel joint model of the sample variability and the differential state of individual transcripts. For each transcript, to minimize the inaccuracy of differential state caused by transcription abundance estimation, we estimate its expression abundance together with the differential state iteratively and enable the differential analysis of weakly expressed transcripts. Simulation analysis demonstrates that the proposed approach has a superior performance over conventional methods (estimating transcription expression first and then identifying differential state), particularly for lowly expressed transcripts. We further apply the proposed approach to a breast cancer RNA-seq data of patients treated by tamoxifen and identified a set of differentially expressed transcripts, providing insights into key signaling pathways associated with breast cancer recurrence.

## Introduction

With the advent of next-generation sequencing technologies, RNA-seq has become a major molecular profiling technique for transcriptome analysis (1–3). The dramatically increased sequencing depth makes it feasible to identify multiple transcripts of the same gene and accurately estimate the abundance (expression) of each, which serves as a base for further differential expression analysis (4–6). While RNA-seq has the advantage of high coverage and fine resolution, there are still many challenges in its analysis like assignment of RNA-seq reads to different transcripts sharing exons; variability of RNA-seq read coverage along genomic loci of transcripts.

Current efforts of gene- or transcript-level differential expression mainly focus on modeling the variability of RNA-seq data between biological samples in the same phenotype group: the between-sample variability (7). Cuffdiff 2 (6) is widely for differential analysis of RNA-seq data at the transcript level. To deal with the within-sample variability, Cuffdiff 2 assumed the same positional bias for transcripts within a certain length range. This assumption, also used in other similar approaches (8), however, is insufficient to account for the complex patterns of within-sample variability observed from data. Moreover, Cuffdiff 2 estimates transcript expression first and detects differentially expressed ones with confident expression, missing many weakly expressed transcripts which can also be differential (9).

In this paper, we develop a novel Bayesian approach for identifying differentially expressed transcripts. This approach is built upon a novel model jointly accounts for both between-sample variability and the differential states of transcripts. Specifically, we use a Poisson-Lognormal model to account for the within-sample variability; a Gamma-Gamma model to analyze the differential expression of transcripts while accounting for the between-sample variability. A Markov Chain Monte Carlo (MCMC) procedure is developed for the joint inference of the model parameters including expression abundance and differential state of each transcript. Simulation studies reveals that our approach has significantly improved the performance in identifying differentially expressed transcripts, especially on transcripts with moderate abundance. We have applied it to a breast cancer RNA-seq dataset and identified a list of transcripts associated with breast cancer recurrence, functioning in diverse breast cancer relevant cellular processes like PI3K/AKT/mTOR signaling and PTEN signaling pathways, which reveals the underlying mechanism of isoforms (transcripts) in driving breast cancer recurrence.

## Methods

There are two types of biological variation in RNA-seq data that affecting accurate quantification of transcript expression: ‘within-sample variability’ and ‘between-sample variability’. The ‘within-sample variability’ refers to the high variance of read counts along the genomic location arising mostly from sequencing bias while ‘between-sample variability’ refers to the variation of read counts (higher than the mean count) among biological replicates or samples. We use a Poisson-Lognormal model to account for the within-sample variability. As different genomic loci may have very different bias patterns, which cannot be well explained by known sources. Using this Poisson-Lognormal model we are able to model different bias patterns along genomic loci at the transcript level. We then use a Gamma-Gamma model to model the transcript expression abundance of multiple samples as well as their differential states between two phenotypes. Specifically, differential states of transcripts are introduced in the Gamma-Gamma model as hidden variables to control the differential abundance of the samples between two phenotypes. Using the above two models jointly, we can model ‘within-sample’ variability and ‘between-sample variability’ simultaneously. As the joint model includes a set of parameters, we use a Markov Chain Monte Carlo (MCMC) procedure including both Gibbs sampling and Metropolis-Hasting sampling to estimate the parameters and the posterior probability of the hidden variable. By virtue of the sampling process, the (marginal) posterior distributions of the parameters and the hidden variable can be estimated (or approximated) by the samples drawn from the MCMC sampling procedure.

### Bayesian model

Let *y*_*t,g,i,j*_ represent the observed counts that fall into the *i*^th^ (1 ≤ *i* ≤ *I*_*g*_) exon region of isoform *t* (1 ≤ *t* ≤ *T*) of gene *g* (1 ≤ *g* ≤ *G*) in sample *j* (1 ≤ *j* ≤ *J*). *T* is the number of isoforms of gene *g* given by the annotation information. *I*_*g*_ is the number of exons in gene *g*. *G* is the total number of genes. *J* = *J*_1_ + *J*_2_ is the total number of samples, where *J*_1_ and *J*_2_ denote the number of samples in phenotype 1 and 2, respectively. Since one gene may have multiple isoforms, *y*_*g,i,j*_, the observed counts in the exon region, is the combination of all potential isoforms, as defined in Eq. (1):

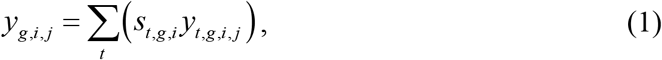

where *s*_*t,g,i*_ is a binary value indicating whether exon *i* is included in isoform *t* of gene *g*. At the isoform level, we use a Poisson-Lognormal regression model to account for the within-sample variability of RNA-seq data. *y*_*t,g,i,j*_ follows a Poisson distribution with mean *γ*_*t,g,i,j*_:

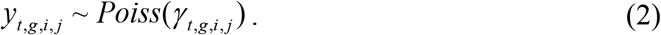

According to the Poisson-Lognormal model (10),

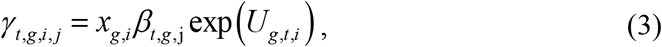

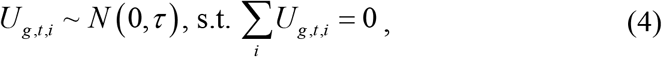

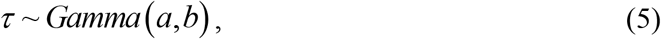

where *β*_*t,g*,j_ is the true expression level of isoform *t* of gene *g* in sample *j*. *x*_*g,i*_ is the length of the *i*^th^ exon weighted by the library size of sample *j*. *U*_*g,t,i*_ is a model parameter representing the within-sample variability (or dispersion) for exon *i* of isoform *t* of gene *g*. Thus, the dispersion of different loci, exons of the isoforms, is modeled by different parameters. Precision parameter *τ* controls the overall degree of within-sample variability.

We use a Gamma-Gamma model (11) to model the expression level *β*_*g,t,j*_ across samples collected from two phenotypes. The differential state, as a hidden variable in this Bayesian model, affects the distribution of *β*_*g,t,j*_ among samples in each of the two phenotypes. *d*_*t,g*_, a binary value, indicates the differential state of isoform *t* of gene g, where *d*_*t,g*_ = 1 means isoform *t* of gene g is differentially expressed; *d*_*t,g*_ = 0, otherwise. Note that the between-sample variability is captured by the Gamma distribution. From the Gamma-Gamma model, the isoform expression level *β*_*g,t,j*_ is given by:

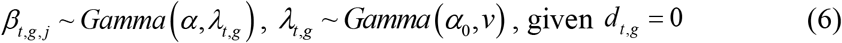

or

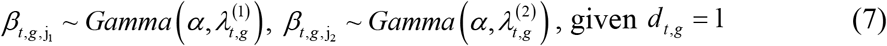

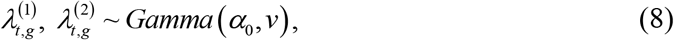

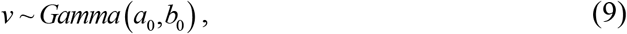

where *α* is the shape parameter; *λ*_*t,g*_ is the rate parameter that depends on differential state *d*_*t,g*_. If *d*_*t,g*_ = 0, 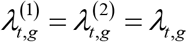; if *d*_*t,g*_ = 1, 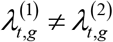. *λ*_*t,g*_ is further assumed to follow a Gamma distribution with shape parameter *α*_0_ and rate parameter *v*. In marked contrast to existing methods like Cuffdiff 2 that uses statistical tests to identify differentially expressed isoforms, the differential states of isoforms are introduced and modeled in the proposed joint model.

### MCMC sampling

We develop a Markov Chain Monte Carlo (MCMC) procedure to estimate transcript differential state (**d**). The likelihood of the observation *y*_*g,i,j*_ given all of the parameters is 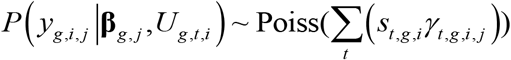. Thus, the conditional (posterior) distributions of the parameters *U*_*g,t,i*_ and *τ* (of the Poisson-Lognormal model) and the parameters **β**, **λ**, *α*, *α*_0_,*v*, and **d** for the Gamma-Gamma model can also be derived (Supplementary Material S1). Then, the MCMC sampling procedure is applied as follows:

***INPUT***: Observed read counts**Y**, library size weighted isoform structure x, number of iterations N

***OUTPUT***: Estimates of all of the parameters and the differential state **d** in the joint Bayesian model

#### Algorithm

**Step 1.** Initialization: each parameter is set an arbitrary value and non-informative prior knowledge is used for the parameters.

**Step 2.** Draw samples iteratively from the conditional distributions of parameters **β**, **U**, ***τ*** (in the Poisson-Lognormal model) and parameters **λ**, *α*, *α*_0_,*v*, and **d** (in the Gamma-Gamma model). Perform the following sampling steps for N iterations:

- Use Gibbs sampling to draw samples of **β**, ***τ***, **λ**, *v* from their conditional distributions that follow standard probability distributions;
- Use Metropolis-Hasting (M-H) sampling to draw samples of **U**, **d**,*α*,*α*_0_ from their conditional distributions in sequence. Since these parameters do not have conjugate priors, M-H sampling is used to approximate their posterior distributions.

**Step 3.** Estimate differential state **d** as well as other parameters **β**, **U**,***τ***, **λ**, *α*, *α*_0_,*v* from the samples, after the burn-in period, generated from the MCMC procedure.

## Results

### Performance evaluation of transcript abundance estimation

Multiple synthetic data sets generated using RNAseqReadSimulator (12) with varying model parameters were generated to compare the proposed Bayesian approach against Cuffdiff 2 (6) and Ballgown (13) for transcript-level differential analysis of RNA-seq data. We first evaluated the accuracy of abundance quantification. Specifically, we used the average correlation of the estimated transcript abundance and the ground-truth transcript expression to assess the accuracy.

First, we varied parameter *τ* to model the overall within-sample variability, with other model parameters set as *α* = 1, *α*_0_ = 0.5, and *ν* = 0.1. In general, the smaller the precision parameter *τ*, the higher the overall within-sample variability. From Fig. 2(a), we can see that, at different *τ*, the average correlations of our estimation were consistently higher than those of Cuffdiff 2 and Ballgown. Note that although RPKM and ENC were calculated by different methods, their correlations to the true expression level across samples were about the same.

Second, we tested the performance of this approach using data with different bias patterns along genomic location. Rather than randomly drawn from *N*(0,*τ*), parameter **U** was designed to follow different bias patterns for different sets of genes. We simulated different patterns (as shown in Fig. 2b) to mimic observed patterns from most real RNA-seq data. Our proposed model is able to capture the bias patterns accurately. Moreover, the performances of our method under different bias patterns are very close, indicating that our model can deal with various bias patterns existing in the same data.

### Performance evaluation of differentially expressed transcript identification

We varied parameter *α* and *α*_0_ in the Gamma-Gamma model to evaluate the performance of our approach on differentially expressed transcript identification. In our model, in general, the less *α* is, the lower abundance of isoform is; the less *α*_0_ is, the more differentially expressed isoforms are. When varying *α*, we set *α*_0_ = 0.5, *ν* = 0.1, *τ* = 1.78; when varying *α*_0_, we set *α* = 1.5, *ν* = 0.1, *τ* = 1.78. For each set of parameters, we generated five replicated experiments and summarized the precision (P), recall (R), and F-measure (F) in Table 1. Results revealed that our method consistently outperformed existing methods. Although Cuffdiff 2 usually has a high precision, it missed many differentially expressed transcripts due the two-step design in their differential analysis. Ballgown could identify more differentially expressed isoforms but with less precision. We can see that our method is very effective when the transcript abundance is generally low (with low *α*) or the transcripts were less differentially expressed (with large *α*_0_). We further evaluated the performance of the competing methods on genes with more than two transcripts (*K*=3, 4, or 5). The experimental results demonstrated that our proposed method outperformed the other methods in multiple scenarios.

**Table 1.** Performance of differential analysis.

| Parameters |  | BayesIso |  |  | Cuffdiff 2 |  |  | Ballgown |  |  |
| --- | --- | --- | --- | --- | --- | --- | --- | --- | --- | --- |
|  |  | P | R | F | P | R | F | P | R | F |
| Abundance control | $\alpha = 1.4$ | 0.919 | 0.543 | **0.683** | 0.894 | 0.22 | 0.353 | 0.814 | 0.362 | 0.499 |
| | $\alpha = 1$ | 0.869 | 0.52 | **0.649** | 0.893 | 0.094 | 0.169 | 0.794 | 0.4 | 0.525 |
| | $\alpha = 0.6$ | 0.824 | 0.446 | **0.577** | 0.796 | 0.114 | 0.199 | 0.655 | 0.258 | 0.367 |
| Differential control | $\alpha_0 = 0.2$ | 0.9 | 0.71 | **0.791** | 0.935 | 0.308 | 0.454 | 0.725 | 0.529 | 0.606 |
| | $\alpha_0 = 0.6$ | 0.898 | 0.513 | **0.652** | 0.842 | 0.239 | 0.37 | 0.743 | 0.393 | 0.511 |
| | $\alpha_0 = 1$ | 0.916 | 0.403 | **0.559** | 0.971 | 0.138 | 0.238 | 0.745 | 0.219 | 0.335 |
| Transcript number control | $K = 3$ | 0.859 | 0.593 | **0.701** | 0.777 | 0.345 | 0.476 | 0.696 | 0.5 | 0.581 |
| | $K = 4$ | 0.809 | 0.538 | **0.642** | 0.69 | 0.261 | 0.374 | 0.657 | 0.321 | 0.389 |
| | $K = 5$ | 0.798 | 0.552 | **0.645** | 0.706 | 0.271 | 0.379 | 0.611 | 0.426 | 0.497 |

### Differentially expressed transcripts associated with breast cancer recurrence

We applied the developed approach to breast cancer data, downloaded from The Cancer Genome Atlas (TCGA) project (12), to identify the differentially expressed transcripts associated with breast cancer recurrence. As cancer data is much noisier than cell line or mouse models, with stronger ‘within-sample’ and ‘between-sample’ variabilities, a strong detection power on moderately differential isoforms would be important for differential analysis of cancer RNA-seq data. In this dataset, there are 93 ER+ patients with 61 patients still alive with the last follow up (survival time longer than 5 years; defined as ‘late/non recurrence’ group) and 32 patients dead within 5 years (defined as ‘early recurrence’ group). 2,299 transcripts of 1,905 genes were identified as differentially expressed between these two groups by our approach. 30% were uniquely identified by our method as compared with the results of applying Cuffdiff 2and Ballgown to the same dataset with default settings. We calculated the SNR of the identified differentially expressed isoforms, with a mean value around −5dB, indicating that most of the identified isoforms are moderately differentially expressed. This low mean SNR value is consistent with the high variability of expression level observed across the samples.

The unique set of differential genes help reveal additional signaling pathways associated with breast cancer recurrence. For example, PIK3R2, a member of the PI3K protein family participating in the regulatory subunit, was detected as significantly down-regulated in the ‘early-recurrence’ group. The loss of expression of PIK3R2 is crucial to the hyperactivation of the PI3K/AKT/mTOR signaling pathway by regulating AKT2. The dysfunction of AKT2 in inhibiting the expression of TSC1 and TSC2 activated the mTOR signaling, as indicated by the over-expression of RPS6KB1, a downstream target of mTOR. The overexpression of TSC2 and RPS6KB1 was further validated by their protein/phosphoprotein expression measured by reverse phase protein array (RPPA) on a subset of the TCGA breast cancer samples, which consists of 45 samples in the ‘late-recurrence’ group and 27 samples in the ‘early-recurrence’ group. Specifically, the expression of transcript NM_001114382, a differentially expressed transcript of gene TSC2, was positively correlated with its phosphoprotein expression at pT1462 (Pearson correlation, *p*-value = 0.02); the expression of transcript NM_001272044, a differentially expressed transcript of gene RPS6KB1, was positively correlated with its phosphoprotein expression at pT389 (p-value = 0.008). Moreover, PTEN signaling, the under-expression of which results in hyperactivation of PI3K/AKT signaling pathway in breast cancer (14,15). Our approached detected SHC1, GRB2 and BCAR1, three critical genes in PTEN signaling.

### Signaling pathways enriched with genes containing differentially expressed transcripts

We conducted a functional enrichment analysis of the identified genes using Ingenuity Pathway Analysis (IPA; http://www.qiagen.com/ingenuity); it tuned out that many of the genes were known to be associated with the following cellular functions: proliferation, cell death, cell migration and several signaling pathways such as Jak-STAT, mTOR, MAPK, and Wnt signaling. We found that many genes associated with the signaling pathways are only differential at the transcript level yet not at gene-level, such as PDPK1, TSC1, TSC2, PIK3R2, AKT2 in the mTOR signaling pathway and HSP90AA1, HSP90AB1 in the PI3K/AKT signaling pathway.

HSP90AA1 and HSP90AB1 are Heat Shock Proteins (HSPs) that play an important role in tumorigenesis (16,17). The overexpression of HSP90AA1 and HSP90AB1 leads to activation of cell viability of tumor cell lines and provides an escape mechanism for cancer cell from apoptosis. Functional analysis using IPA has shown that the down-regulation of HSP90AB1 leads to activation of cell death of immune cells (18). While HSP90AA1 has two isoforms from alternative splicing, only NM_005348 (RefSeq_id) was overexpressed in the ‘early-recurrence’ group. HSP90AB1 has five isoforms, among which NM_007355 was detected as overexpressed in the ‘early-recurrence’ group whereas NM_001271971 was overexpressed in the ‘late/non recurrence’ group. These findings suggest that the change in different expression pattern of the transcripts might contribute to different functions of cancer development.

## Discussion

We have developed a Bayesian approach for the identification of differentially expressed transcript. It is a fully probabilistic approach that estimates the differential states and other model parameters like transcript abundance iteratively. Differential analysis benefits from the accurate estimation of transcript expression, especially when the ‘the between-sample variability’ of transcript expressions is high. Thus, transcript expression estimation and differential state inference are tightly coupled in our framework. Not only for differential analysis, the accurate abundance estimation at transcript level also provides high quality input data for regulatory network analysis (19,20), for validating target genes of specific upstream regulators (21)(22) or for identifying rewiring of signal pathways in different conditions (23).

However, it is a non-trivial task to model the sequencing bias for RNA-seq data analysis. The bias patterns are complicated and cannot be well explained by known sources. We used a flexible model to account for the bias independent of any particular pattern. Certain bias patterns (such as bias to the 3’ end, or high in the middle) occur more frequently than others. Moreover, we also observed that the bias patterns can be affected by the expression level. In the future work, we will incorporate certain bias patterns as prior knowledge into the model, which can help estimate the bias pattern of some transcripts more accurately.

